# Therapeutic Challenges of Multidrug Resistant *Acinetobacter baumannii* in Eastern Africa: Systematic Review

**DOI:** 10.1101/558312

**Authors:** Alene Geteneh, Assalif Demissew, Alemale Adimas, Derbie Alemu, Lencho Girma

## Abstract

*Acinetobacter baumannii*, an opportunistic gram negative bacterium, is known to emerge as a major health threat in Eastern Africa. Clinical isolates exhibit resistance to carbapenems and most or all available antibiotics. This review is intended to present concerns about resistance and therapeutic challenges of multi drug resistance *Acinetobacter baumannii* in Eastern Africa. Data was obtained from PubMed and Google scholar, and from free goggle access and web Medline for facts about *Acinetobacter baumannii* and its resistance pattern. Moreover, Preferred Reporting Items for Systematic Reviews and Meta-Analysis (PRISMA) flow chart was used to guide the selection of study materials. Total of 98 articles identified, 13 fit the criteria and were included for the final analysis. In East Africa the overall prevalence of *Acinetobacter baumannii* was 4.95%, while the overall rate of multi drug resistance, carbapenem and pan resistance was 87.3%, 64.8% and 25.2% respectively. Colistin resurges as potential therapeutic options to overcome the lack of new antibiotic treatment of *Acinetobacter baumannii.* There needs a collaborative effort in researches targeted for *Acinetobacter baumannii* treatment and respond for call of “Research and Development of new antibiotics” to control its damning impact.

## INTRODUCTION

*Acinetobacter baumannii* is an opportunistic bacterial pathogen primarily associated with hospital-acquired infections. Genus Acinetobacter is a strictly aerobic, non-fastidious, non-motile, oxidase-negative, non-lactose fermenter, catalase positive, gram-negative coccobacilli that is found as ubiquitous saprophytes (1–3). Unlike *Acinetobacter baumannii*, *A. lwoffi* and *A. haemolyticus* are non-glucose oxidizing species in this genus with less common role in disease causation (4). While, *A. baumannii* takes the lions share because of its increasing involvement in a number of severe infections and outbreaks occurring in clinical settings with an intrinsic or acquired resistance to multiple classes of antibiotics(5, 6).

*Acinetobacter baumannii* has come to light as a major threat to the global health, with a rapid expansion of resistance to carbapenems and most or all available antibiotics (7, 8).

The growth of resistance to at least one agent in three or more classes of antibiotics (aminoglycosides, carbapenems, cephalosporins, beta-lactams/beta-lactamase inhibitor, quinolones etc.) defined the development of multidrug-resistant *Acinetobacter baumannii* (MRAB) traits (9–12). This notable success is granted by its strange capacity to develop or acquire antibiotic resistance determinants and by its ability to adapt and survive for long periods in the hospital environment (13, 14).

*Acinetobacter baumannii* appears well apt for genetic exchange and are among a unique class of gram-negative bacteria that are described as “naturally transformable”(15, 16). Antibiotic resistance in *A. baumannii* is due to enzymatic degradation, modification of targets, and active efflux of drugs collectively with formation of biofilms (16, 17). Although outbreaks are typically clonal, horizontal-gene transfer has an important role in broadcasting of antibiotic resistance to *A. baumannii* and pass on *bla* _NDM_ gene to other *Enterobacteriaceae* (18–20).

Although, there is no organized data in developing countries who administered broad spectrum antibiotics empirically and had limited facility for isolation and susceptibility testing of bacteria for patient care, there could have high distribution of MRAB even resistant to carbapenems. *Acinetobacter baumannii*, being resistant for multiple antibiotics also results in high case-fatality rate 38% in Kenya(21) and 32% in South Africa(22)) due to an overwhelming systemic infection and lack of treatment options(23).

Updated realization is very essential for the prevention of MRAB infections and to cope up with therapeutic challenges, thus we had included 13 recent articles of East Africa (published from 2013-2017, from Uganda, Tanzania, Sudan and Ethiopia). The objectives of this review were to consolidate existing data about the resistance and therapeutic challenges of MRAB, and to show its burden in Eastern Africa.

## METHODOLOGY

Here in this update we have searched for articles using mesh terms like *Acinetobacter baumannii*, multidrug resistant *Acinetobacter baumannii*, “ *(Acinetobacter baumannii)* AND multidrug resistant AND Uganda*“* (multidrug [All Fields] AND resistant [All Fields]) AND (“*Acinetobacter baumannii*” [MeSH Terms] OR (“*Acinetobacter*” [All Fields] AND “*baumannii*” [All Fields]) OR “*Acinetobacter baumannii*” [All Fields]) AND (“Uganda” [MeSH Terms] OR “Uganda” [All Fields]) AND (“2013/01/01” [PubDate]: “2017/12/31” [PubDate]), (multidrug [All Fields] AND resistant [All Fields]) AND (“*Acinetobacter baumannii*” [MeSH Terms] OR (“*Acinetobacter*” [All Fields] AND “*baumannii*” [All Fields]) OR “*Acinetobacter baumannii*” [All Fields])) AND (“Ethiopia” [MeSH Terms] OR “Ethiopia” [All Fields]) AND (“2013/01/01” [PubDate]: “2017/12/31”[PubDate]) and similarly for other East African countries from PubMed, and Google scholar, free goggle access and web Medline for facts about *Acinetobacter baumannii* and its resistance pattern. Moreover, PRISMA flow chart was used to guide the selection of study materials.

### Selection criteria

Any sort of article who mentioned the prevalence of *Acinetobacter baumannii*, described patterns of resistance for commonly used antibiotics (resistance to multiple antibiotic including carbapenem and colistin) and being in an Eastern African region were included.

### Inclusion (Eligibility) and exclusion criteria

In this review we have been looking for studies (both prospective and retrospective), not more than 5 years and study theme similar to the review questions were included, and studies done on non-clinical samples, review articles, articles without pdf access and papers describing isolates outside Eastern Africa were excluded.

### Data extraction

Data extraction was done manually using Microsoft Excel 2010. Information extracted included article information (country, year of publication, number of participants enrolled, number of participants positive for *A. baumannii*), study design (sample size and types of specimens used), and antimicrobial resistance data.

### Data analysis

Based on the review questions sought, we compile data on Microsoft excel 2010 and transported to STATA for use to calculate proportion, ratio and mean resistance to highlight the pattern of antibiotic resistance among isolates in eastern Africa context.

### RESULTS

Of the 98 articles identified, 13 fit the criteria and were included for the final analysis (Fig.1). Majority of the articles were from Uganda (5) and Ethiopia (4). As shown in the table the overall prevalence of *A. baumannii* was 4.95 % [95% CI; 0.8 −9%], while the overall proportional rate of MDR, carbapenem and pan resistance was 87.3% [95%CI; 78.2-96.5%], 64.8% [95% CI; 27.7-100%] and 25.2% [95% CI; 7.3-43%] respectively.

**Figure 1:**
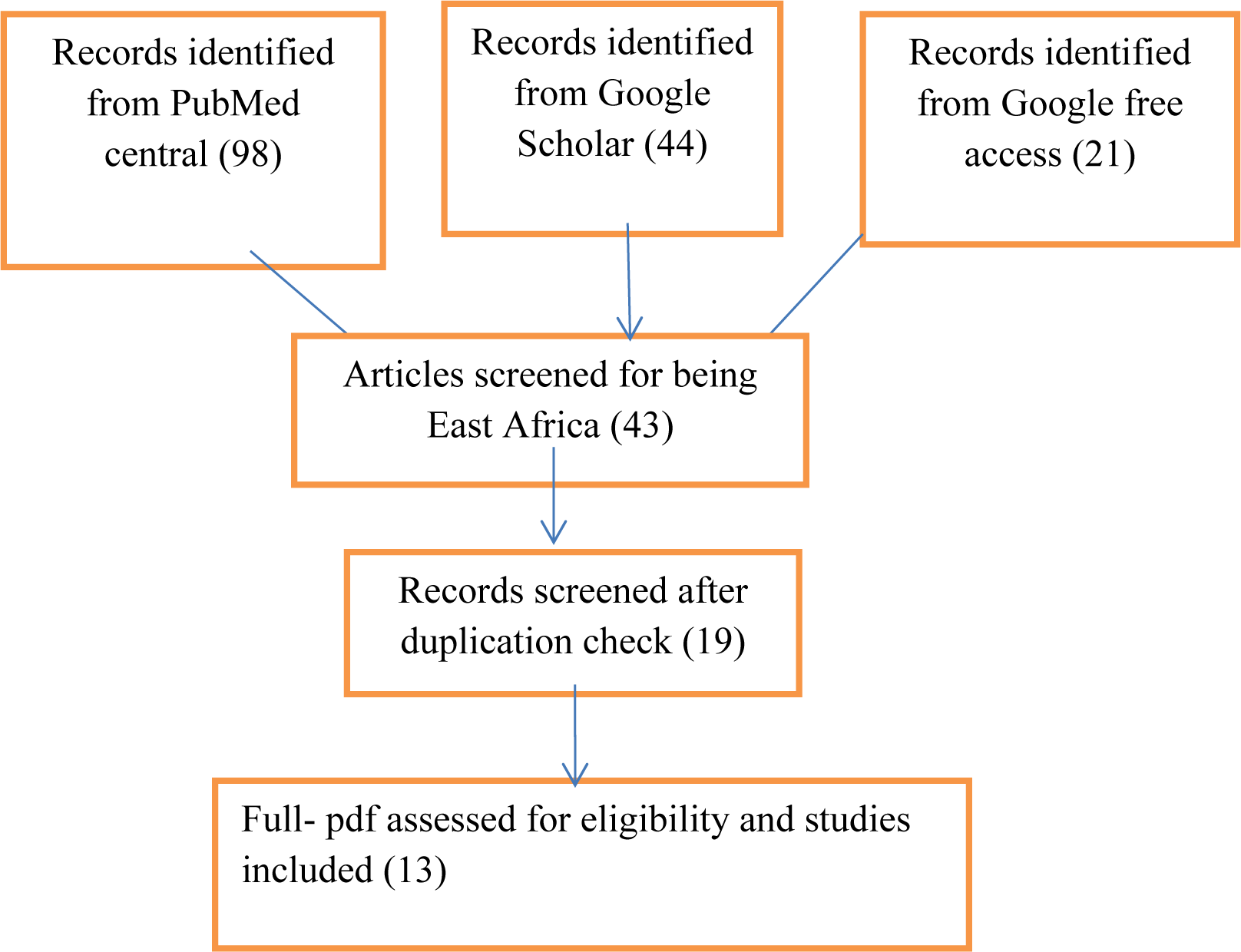
PRISMA Diagram of the article selection procedure for articles published from 2013 -2017 * Preferred Reporting Items for Systematic Reviews and Meta-Analysis

The average number of patients included were 710 (ranging from a minimum of 100 to a maximum of 2899), of them nearly 5% of patients enrolled were positive for *Acinetobacter baumannii.* On other hand, the mean proportional rate of resistance for multiple antibiotics was so high, provide 87.3% (nearly 31 out of 36) (Table 2).

**Table 1:** Summary of selected articles in Eastern Africa for multi-drug resistance *Acinetobacter baumannii* (2013–2017)

|  | Study design | Country | Year of publication | Patients enrolled | Patients positive | No. of MDRAB | CRAB | Pan resistance | Reference |
| --- | --- | --- | --- | --- | --- | --- | --- | --- | --- |

|  |  |  |  |  | for AB |  |  |  |  |
| --- | --- | --- | --- | --- | --- | --- | --- | --- | --- |
| 1. | Cross-sectional | Ethiopia | 2013 | 322 | 7 | 4 | 0 | 0 | (24) |
| 2. | Cross-sectional | Uganda | 2013 | 314 | 52 | 52 | 2 | 2 | (25) |
| 3. | Cross-sectional | Tanzania | 2014 | 100 | 15 | 15 | 6 | 0 | (26) |
| 4. | Cross-sectional | Tanzania | 2015 | 469 | 6 | 0 | 0 | 0 | (27) |
| 5. | Cross-sectional | Sudan | 2015 | 2899 | 275 | 253 | 244 | 102 | (28) |
| 6. | Cross-sectional | Uganda | 2016 | 869 | 29 | 18 | 9 | 0 | (29) |
| 7. | Cross-sectional | Ethiopia | 2016 | 107 | 23 | 22 | 19 | 8 | (30) |
| 8. | Cross-sectional | Uganda | 2017 | 111 | 11 | 7 | 7 | 1 | (31) |
| 9. | Cross-sectional | Uganda | 2017 | 736 | 7 | 7 | 1 | 0 | (32) |
| 10. | Cross-sectional | Tanzania | 2017 | 575 | 5 | 1 | 0 | 0 | (33) |
| 11. | Cohort | Uganda | 2017 | 1786 | 11 | 10 | 0 | 0 | (34) |
| 12. | Cross-sectional | Ethiopia | 2017 | 224 | 14 | 8 | 8 | 2 | (23) |
| 13. | Cross-sectional | Ethiopia | 2017 | 712 | 2 | 2 | 0 | 0 | (35) |
|  | Total |  |  | 9224 | 457 | 399 | 296 | 115 |  |

**Table 2:**
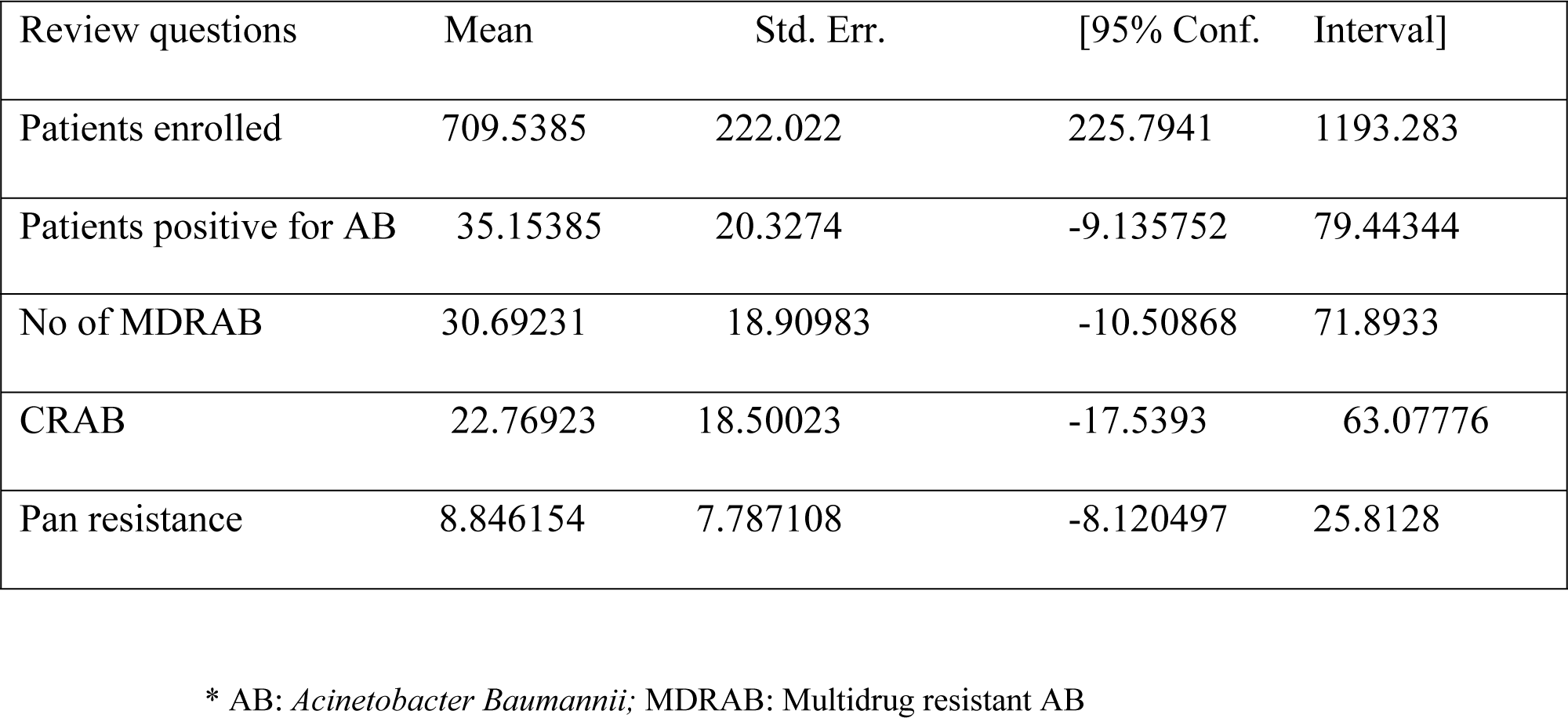
Overall mean estimate of *Acinetobacter baumannii* in East Africa (2013–2017)

Considering Eastern Africa as one sample population at a snapshot of review period, we tried to see ups and downs in incidence, and resistance for multiple antibiotics, carbapenems and all available treatments options as well. Therefore, the overall positivity rate in patients screened for *Acinetobacter Baumannii* was shown to decrease from 2013 to 2017 (Figure 2).

**Figure 2:**
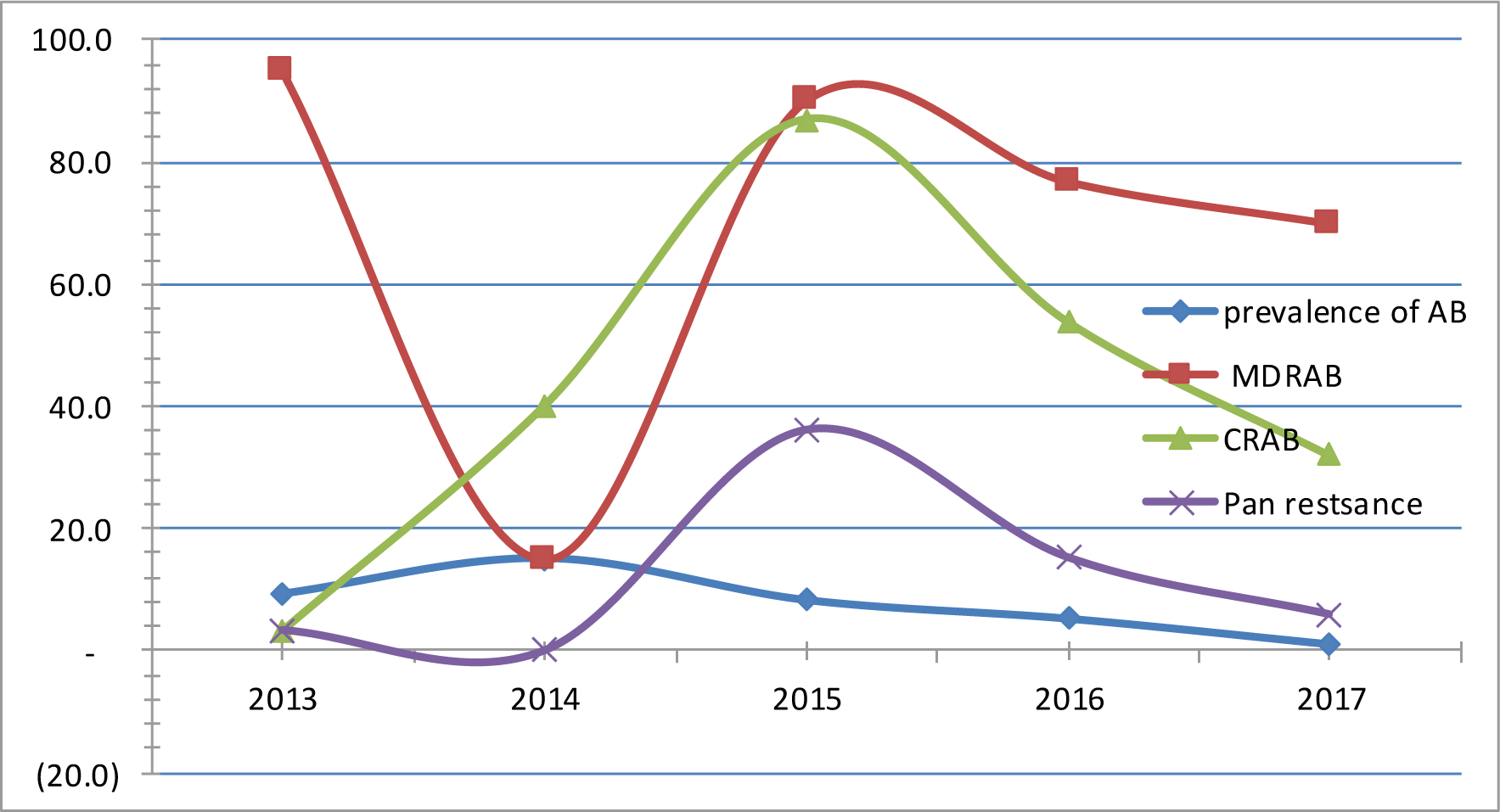
Overall prevalence of AB, MDRAB, CRAB and pan resistance with year of publication * AB: *Acinetobacter Baumannii;* MDRAB: Multidrug resistant AB; CRAB: Carbapenem resistance

The amalgamated prevalence and pan-resistance rate downs to nullity, while the resistance rate for multiple drugs was relatively stable during 2016 to 2017(Figure 2). There were variation in state of decrement in prevalence, proportional rate of MRAB, CRAB and pan-resistance according to an individual country data of Ethiopia and Uganda (Figure 3 and figure 4).

**Figure 3:**
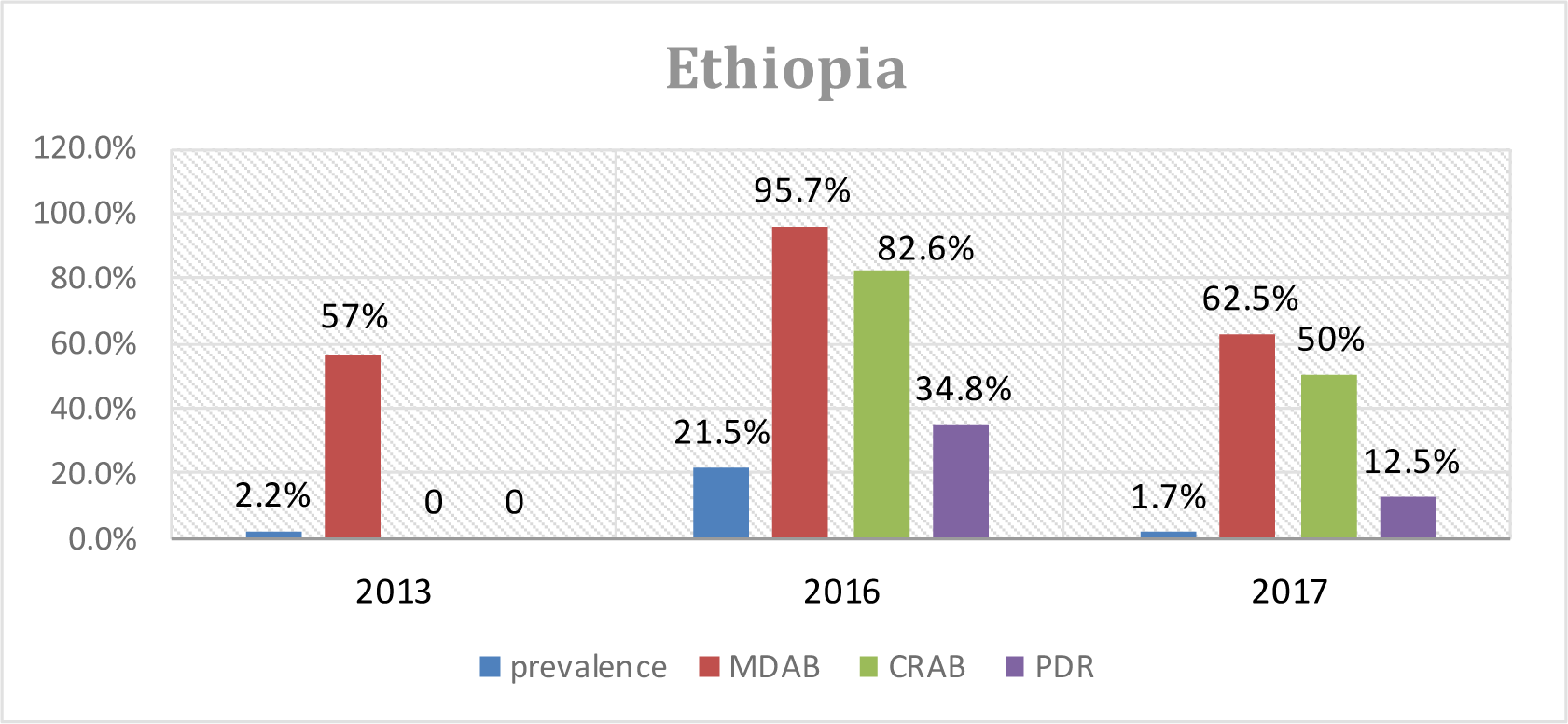
Prevalence and proportional rate of resistance in isolates in Ethiopia, 2013-2017

**Figure 4:**
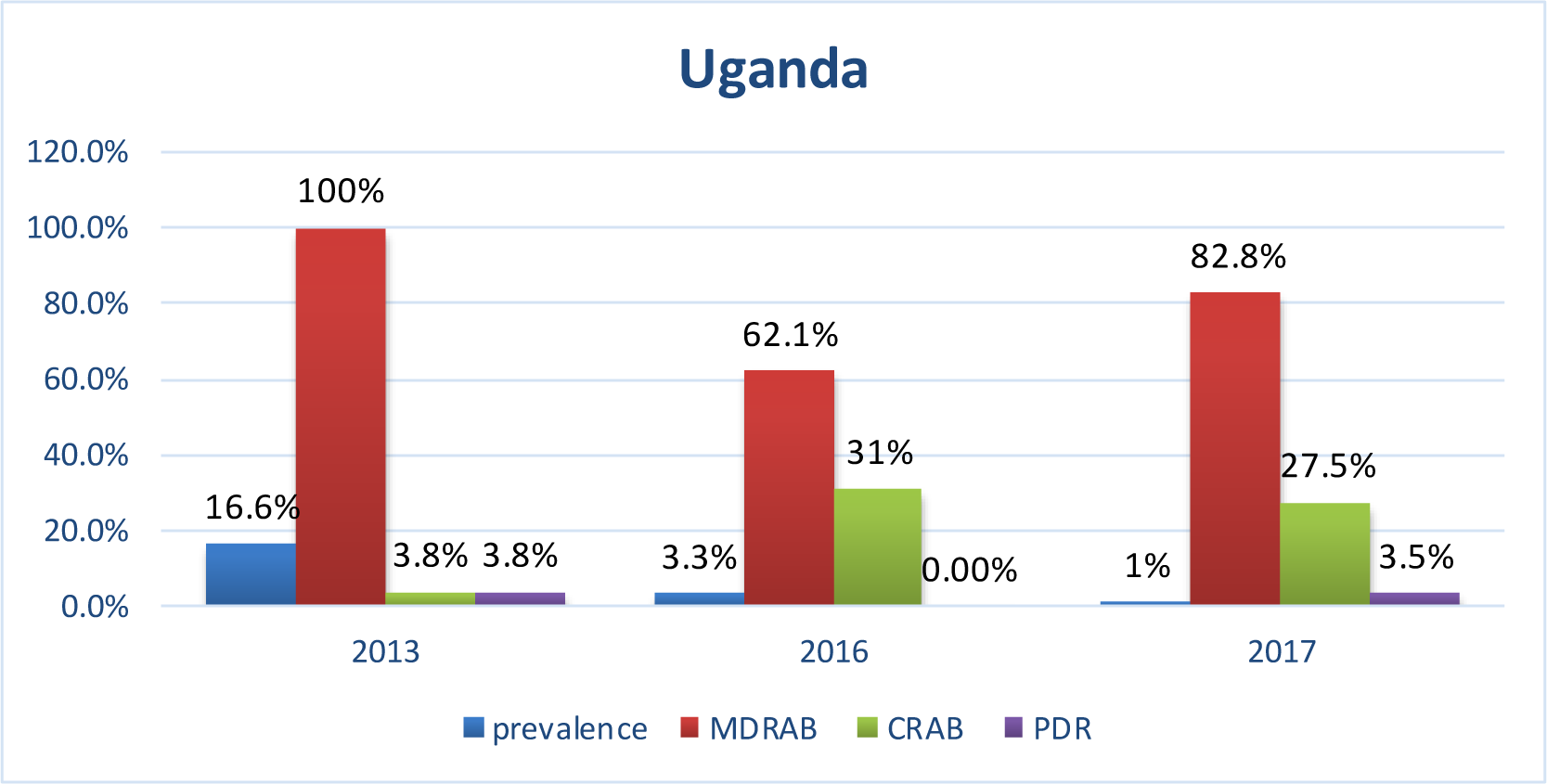
*Acinetobacter baumannii* isolates incidence rate and proportional rate of resistance in Uganda, 2013-2017

All of the isolates in the revision were only from clinical specimens such as blood, urine, wound swabs, sputum, tracheal swabs and CSF etc., to show its detectability with culture (all the articles use culture for isolation).

## DISCUSSION

Multidrug-resistant *Acinetobacter baumannii* infections posed a serious health trouble all over the world (36, 37). Even though, the overall prevalence (4.95%) was not considerably high, our review assured that resistance to multiple antibiotics was also a great concern for Eastern Africa (87.3%). Similarly carbapenem-resistant *Acinetobacter baumannii* (CRAB) were found the most important group of actors in the ongoing antibiotic resistance crisis, i.e. proportionally 64.8% were resistant to carbapenems. This incidence is nearly comparable with a study finding in Egypt (70%) (38).

World health organization (WHO) has prioritized *Acinetobacter baumannii* as a “critical” pathogen for Research and Development of new antibiotics; and old antibiotics such as fosfomycin and polymyxins was got considerations as potential treatment options to prevail over the lack of new antibiotics (13, 39–41). This was true for East African countries, because proportionally 87.3% of the isolates want to be treated with carbapenems, but the pragmatic concern is 64.8% of them were resistant for these drugs putting its user-friendliness into consideration.

Our findings were higher as compared to study of India, which was 42.3% and 6.7% carbapenem and pan resistance respectively (42). Similarly, extensively drug resistant (resistance to at least one agent in all but two or fewer antimicrobial classes (12)(XDR)) phenotypes exhibited via carbapenem resistance were nearly equivalent to Pakistan (65.6%), but higher prevalence of MDR (96.7%)(43), relative to this review (87.3%). This difference might due to variability in socio-demography, specimen used and drug resistance pattern was checked from the already isolated strains, not from direct patient samples of Pakistan and India.

Generally, there has been a relative decrement from 2013-2017 in prevalence, resistance to multiple antibiotics, resistance to carbapenem and pan resistance as shown in figure 2. But, the progress of those MRAB strains to XDR phenotypes is the other worrying issue. In our case scenario, 74.2% MDRAB isolates were XDR phenotypes, exhibited by carbapenem resistance (13) making the deal particularly challengeable for clinicians. The growing incidence of XDRAB infections, therefore quests for salvage therapy with colistin, amikacin, or tigecycline.

The specific country based data has also shown the proportional decrement in prevalence of *A. baumannii* in Uganda from 2013, 2016 and 2017(16.56%, 3.34% and 1.1% respectively), while in Ethiopia fluctuated incidence have occurred (2.17%, 21.5% and 1.7%) with respected years of publications. Similarly the proportional rate of resistance to carbapenem and pan drugs seems to decline in both Ethiopia and Uganda with time, as shown in figures 3 and 4.

Overall, the possible justification for reduction in prevalence and resistance in Eastern Africa may be due to research limitation to figure out the real burden in study areas, or the articles included with low number of study subjects may not represent the actual situation in the area, or use of strategies that may restrict broad spectrum antibiotics administration, the raise in awareness about multidrug resistance, delivery of guidelines to an international level, presentation of strategies in health care facilities and involvement of communities, and implementation of comprehensive control plans.

On the other hand, the average figure (4.95%) seems to represent a small prevalence, but the high multidrug-resistance pattern poses difficult therapeutic challenges. For example, the mortality rate due to this super-bug was much higher in South Africa (44.6%) (44), Kenya (38%) (21), although the articles which we analyzed did not specify *A. baumannii* related deaths. Those patients infected with carbapenem resistant strains have a 35.2% proportional chance of cure with these honored drugs and seeks for easily inaccessible drug colistin and resistant to 25.2% of those who had the chance in the poor nations of Eastern Africa.

The surgical site infections due *A. baumannii* in Ethiopia shown that 34.8% of the isolates were pan-resistant, unable to treat with locally available treatment options leading for unnoticed death (30). Similarly, 16.3% of isolates from the operation theatre and delivery rooms were responsible for untreatable infections to patients attending hospitals of Ethiopia (45). Whereas, in Uganda 55% of hospital environment isolates were resistant for carbapenem looking for the unavailable and 25.2% proportionally resistant drug, colistin (29).

Inappropriate empiric treatment administration results in resistance to multiple antibiotics and this has been associated with 2 to 4 fold increased risk of mortality (10). The high (87.3%) MDR proportion with other comorbidities and short handiness to better treatment options may let for a considerable death in Eastern Africa that we may not noticed. *A. baumannii* infections was known to be nosocomial, but recently there were reports of high prevalence community acquired infections, exemplified by the high prevalence of *A. baumannii* among head and body lice of Ethiopia (46), and community acquired blood stream infections of Tanzania (27).

## CONCLUSION AND RECOMMENDATIONS

A considerably rising rates of MDR as well as XDR *A. baumannii* has distributed throughout the study countries, assuring its major threat to the health of eastern Africa. Old antibiotics such as colistin resurge as potential therapeutic options to overcome the lack of new antibiotics, but the simultaneous emergence of colistin-resistant strains initiate necessity of new advances for control. Which in turn, quest researchers to respond to the call for “Research and Development of new antibiotics” to this super-bug.

## Declarations

The authors declare that there is no conflict of interest regarding the publication of this paper.

---

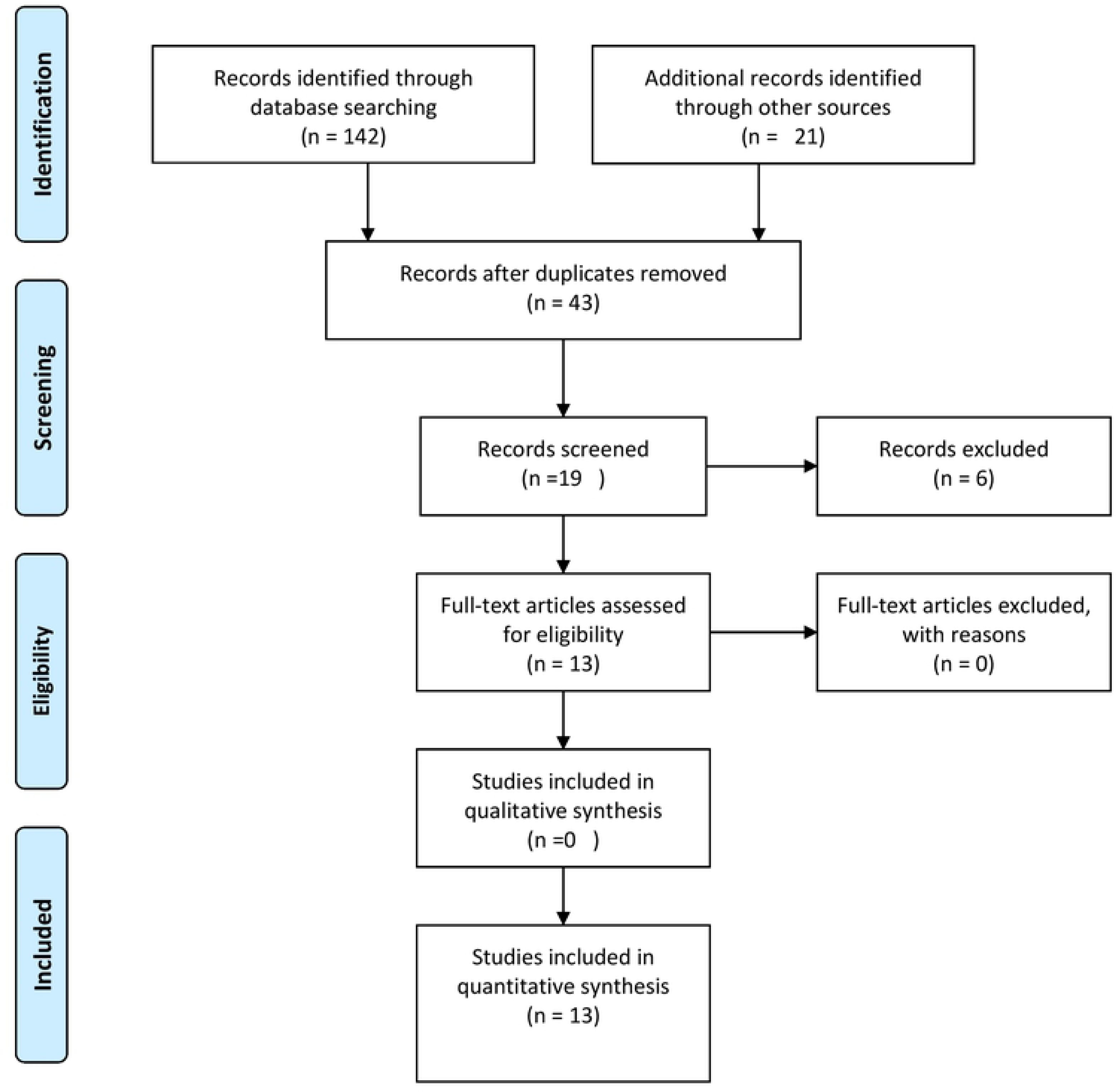
PRISMA Flow Diagram.

